# Music Reveals Medial Prefrontal Cortex Sensitive Period in Childhood

**DOI:** 10.1101/412007

**Authors:** Laurel J. Gabard-Durnam, Takao K. Hensch, Nim Tottenham

## Abstract

An outstanding issue in our understanding of human brain development is whether sensitive periods exist for higher-order processes (e.g., emotion regulation) that depend on the prefrontal cortex. Evidence from rodent models suggests that there is a sensitive period before puberty when acoustic stimuli, like music, shape medial prefrontal cortex (mPFC) responses that regulate affect in the context of acute stress in adulthood. The present study examined whether a homologous sensitive period for the mPFC occurs during human childhood. In the context of acute stress, adult behavioral preferences were observed only for music experienced during childhood, not preschool or adolescent periods. Childhood music increased mPFC activation and modulated connectivity with the amygdala, which was associated with enhanced emotion regulation and lowered autonomic arousal. Moreover, the timing of this sensitive period could be moved by early-life stress. These findings indicate that childhood is a sensitive period for mPFC encoding of regulatory stimuli.

Sensitive periods are developmental moments of heightened neuroplasticity when experiences shape brain function and behavior with lasting effects (*1*). They are fundamental to human cortical ontogeny, and yet, the timing and nature of sensitive periods for human prefrontal cortex functions remain unknown (*2*–*8*). In the rodent, Yang and colleagues have shown that the medial prefrontal cortex (mPFC) exhibits a sensitive period during the prepubertal juvenile period, with heightened responsivity to complex auditory stimuli (i.e., music) (*9*). Specifically, initial exposure to music during the open (juvenile) sensitive period or a pharmacologically re-opened sensitive period in adulthood was followed by that music uniquely producing a behavioral preference in adulthood, increasing mPFC activity, and reducing anxiety-like behavior. This finding and others converging on the same prepubertal period of plasticity in the rodent (*9*, *10*) may have important implications for human development; the complementary prepubertal period in humans (i.e., school-age childhood) also exhibits developmentally-unique mPFC circuitry phenotypes (*11*–*17*) that make childhood a strong candidate for a human mPFC sensitive period. Here, we used Billboard music chart data to identify age-specific exposures to pop songs to test whether music shapes human mPFC responses during a childhood sensitive period as in the rodent. To parallel the approach used in the rodent, we examined whether (i) music experienced during childhood uniquely produced a behavioral preference under stress, (ii) childhood music enhanced emotion regulation behaviorally and physiologically, (iii) mPFC activity was enhanced by childhood music, and mPFC circuitry mediated emotion regulation benefits of childhood music, and (iv) whether the timing of this putative sensitive period could be shifted by early adversity.

## Body of text

Adults born in the USA with developmental exposure to pop music (childhood-exposed group) and adults who recently immigrated to the USA with no exposure to American pop music in childhood (control group) were administered an acute lab stressor (several blocks of a speeded math test). Between stressor blocks, they were given rest periods during which they made behavioral choices between radio stations that played U.S. pop song clips (Figure 1a and b). One station played U.S. pop songs coinciding with middle childhood, the putative mPFC sensitive period, while a second station played U.S. pop songs from late adolescence (post-putative sensitive period) (Figure 1 left panel). A second replication sample assessed the specificity of the childhood period by including music from a preschool period.

**Figure 1.**
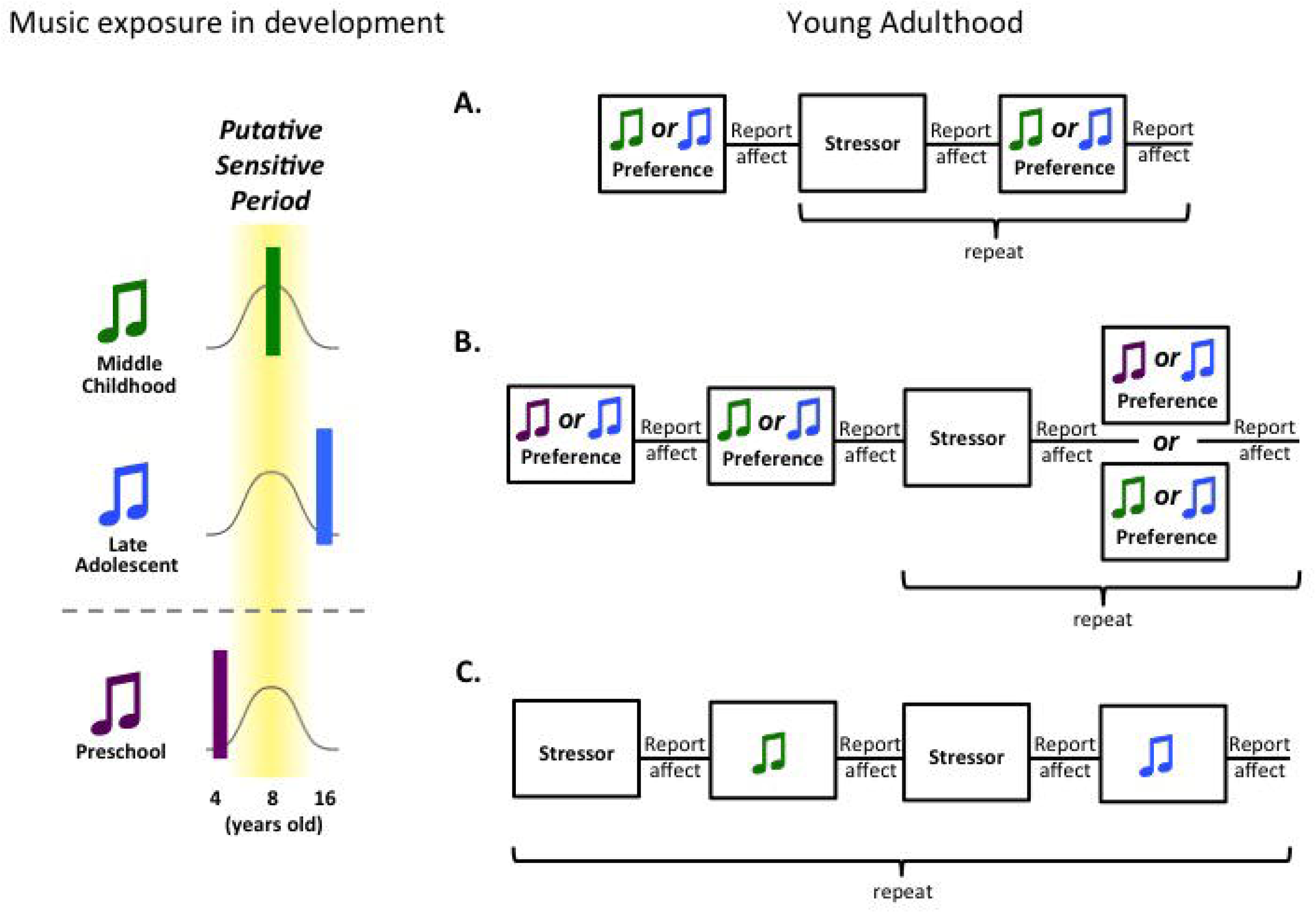
Study schematic. Left panel: Pop music played during different developmental periods that correspond to stages of the putative sensitive period: during the sensitive period (middle childhood music), after the sensitive period (late adolescent music), and before the sensitive period (preschool). Study paradigms (A, B, C) were completed in young adulthood. A. Behavioral preference paradigm: participants chose between two radio stations, one playing childhood songs and the other playing adolescent songs, interwoven with stressor blocks. B. Behavioral preference paradigm including preschool songs in addition to childhood and adolescent songs. C. Functional magnetic resonance imaging paradigm: participants listened to songs alternating between the childhood and adolescent periods interwoven with stressor blocks.

### Stress Induces a Behavioral Preference for Childhood Music

Prior to the stress paradigm, individuals’ baseline behavioral preferences for each music period (percent dwell time listening to that station) were established. Following stress, behavioral preferences for the music periods differed according to developmental exposure levels (F_(2, 91)_ = 6.81, p = 0.002, n = 95; Figure 2A). Childhood-exposed adults increased their behavioral preference for childhood music following stress. For participants who reported a higher degree of exposure in childhood (i.e., median split), the preference for childhood music became the majority choice after post-stressor (mean 58.4% dwell time, CI_L95_ 51.5% - CI_U95_ 65.3%). Childhood-exposed adults with low childhood exposure and the control group maintained preferences for adolescent music across baseline and stress contexts (low exposure dwell time: baseline mean 31.4%, CI_L95_ 23.0% - CI_U95_ 39.7%; post-stressor mean 37.2%, CI_L95_ 28.7% - CI_U95_45.6%; control group: baseline mean 31.8%, CI_L95_ 24.9% - CI_U95_ 38.9%; post-stressor mean 24.1%, CI_L95_ 17.0% - CI_U95_ 31.3%). The childhood-exposed group did not report explicitly liking one music period’s songs more than the other (t(56) = -0.174, p = 0.863, n = 57). A secondary sample of USA-born adults completed the paradigm with earlier preschool music (pre-putative sensitive period, Figure 1 left panel) to test the temporal specificity of the childhood music preference (Figure 1b). Childhood-exposed adults showed a relative preference for childhood compared to preschool music (t(19) = 2.54 p = 0.01, n = 21), and they showed no behavioral preference for preschool relative to adolescent music (preschool music dwell time: mean 38.2%, CI_L95_ 31.5% - CI_U95_ 48.4%; Figure 2B), indicating the behavioral preference under stress is specific to childhood, and does not reflect a general familiarity or recency effect.

**Figure 2.**
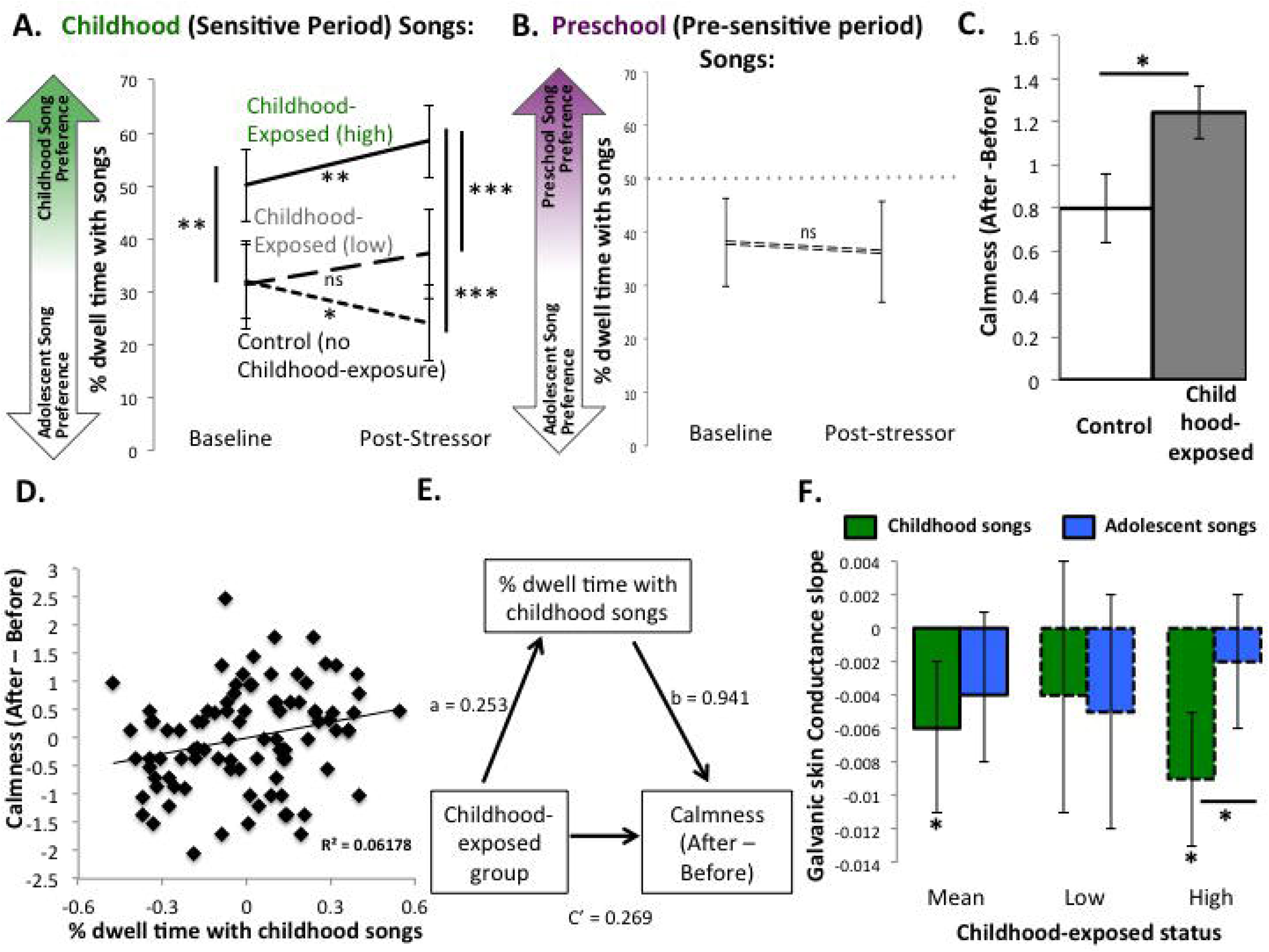
Adult preference is specific to middle childhood music, which has anxiolytic effects under stress. A. Behavioral preferences (percent dwell time) for childhood songs relative to adolescent songs (y axis) before and after the stressor (x axis) for the control (no childhood-exposure) group and the childhood-exposed group. B. No behavioral preference for preschool songs relative to adolescent songs. C. The childhood-exposed group reports greater increase in calmness than the control group after the music blocks. D. Association between dwell time with the childhood songs during music blocks and reported calmness after music blocks. E. Bootstrapped mediation model of childhood-exposure group (childhood-exposed or control) modulating reported emotion regulation through preference (% dwell time) for childhood music post-stressor. F. Galvanic skin conductance slope (change in arousal) during the childhood songs and adolescent songs within the childhood-exposed group as a function of exposure level during development. * p < 0.05. ** p < 0.01. *** p < 0.001. Error bars represent 95% confidence intervals around the mean.

### Childhood Music Regulates Negative Emotion Behaviorally and Physiologically

The effect of music behavioral preferences on emotion regulation under stress was assessed through self-report and physiological recordings. Following the music blocks in the behavioral stress paradigm, the childhood-exposed group reported feeling calmer than the control group (b = .446, t = 2.13, p = 0.036, n = 93, Figure 2C). This association between childhood exposure and calmness was mediated by the amount of dwell time participants spent with childhood music (indirect effect: 0.177 (SE 0.10), CI_L95_ 0.006 – CI_U95_ 0.407, n = 93; Figure 2D and E). The same mediation was observed when examining childhood exposure variation within only the childhood-exposed group as well (indirect effect: 0.193 (SE 0.10), CI_L95_ 0.034 – CI_U95_ 0.447, n = 59).

Physiological differences in emotion regulation as a result of behavioral preferences were assessed using galvanic skin conductance (GSC) slope within the childhood-exposed group. Steeper declines in GSC were observed when participants listened to childhood relative to adolescent music, an effect largely driven by those with higher childhood exposure (stimuli x childhood exposure F_(1, 43)_ = 5.849, p = 0.02, n = 47; Figure 2F). Adolescent music did not result in any significant declines in GSC slope (all confidence intervals included 0). That is, within individuals developmentally exposed to both music periods, only childhood music resulted in a significant autonomic arousal decrease in adulthood.

### Medial prefrontal Cortex Response Reflects Childhood Music Exposure

A subset of participants underwent functional magnetic resonance imaging (fMRI) during a similar math stressor paradigm, interwoven with blocks alternating between childhood and adolescent song clips (Figure 1C). There were no group differences in fMRI reactivity during the math stressor blocks (voxelwise uncorrected p < 0.005, FWE alpha < 0.01). A whole-brain analysis revealed an Exposure Group (childhood-exposed vs. control) X Stimulus Presentation (childhood vs. adolescent) interaction in the mPFC (voxelwise uncorrected p < 0.005, FWE alpha < 0.01). Post hoc tests revealed that supragenual anterior cingulate cortex (supraACC) (center of mass: x,y,z = 3, 32, 21, Figure 3A) showed unique activation to childhood music in the childhood-exposed group (exposed group mean childhood music>baseline reactivity = 0.122, CI_L95_ 0.0259 – CI_U95_ 0.218, t = 2.767, p = 0.017, n = 13; all other reactivity = p > 0.1; Figure 3A). SupraACC responsivity to childhood music was not associated with later-life exposures or reported liking of childhood music as an adult (exposure and liking p > 0.1), yet it predicted subsequent affect, such that greater supraACC reactivity predicted decreased reported negative affect after the music (b: -0.249, t = -2.352, p = 0.030; Figure 3B). For complete fMRI results, see Supplemental Results.

**Figure 3.**
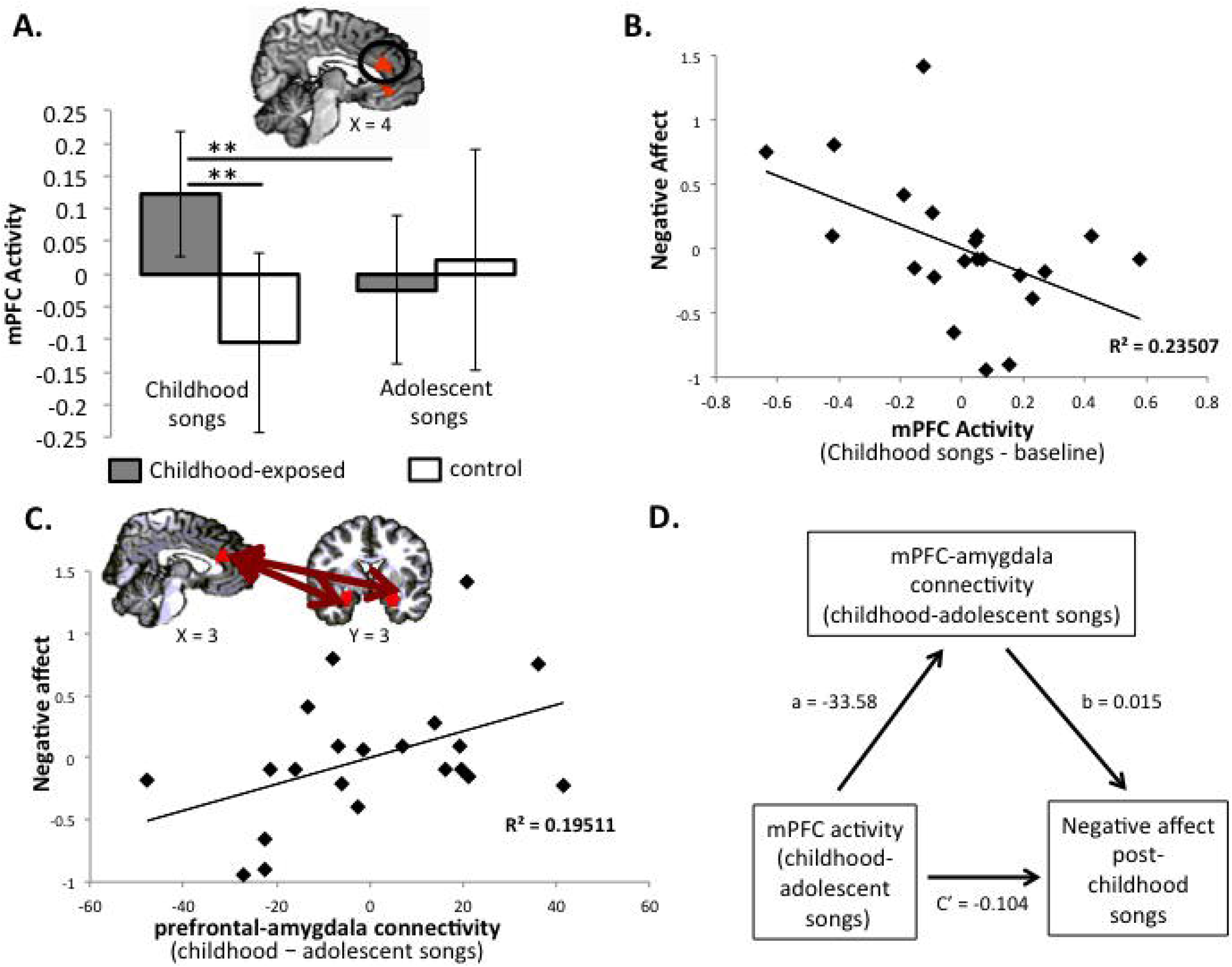
Medial pre-frontal cortex responds to childhood music and is associated with anxiolytic effects of childhood music. A. Whole-brain functional magnetic resonance imaging (fMRI) analysis showing medial prefrontal cortex region (in black circle) differentiates childhood relative to adolescent songs between childhood-exposed and control (no childhood-exposure) groups. P < 0.005, alpha < 0.01, corrected. fMRI parameter estimates are shown for each condition (versus implicit baseline). B: Medial prefrontal cortex (mPFC) reactivity to childhood songs (versus implicit baseline) (x-axis) predicts self-report of negative affect after listening to childhood songs (y-axis). C. Functional connectivity between the medial prefrontal cortex (mPFC) and bilateral amygdala during childhood relative to adolescent songs (x axis) predicts self-report of negative affect after listening (y-axis). D. Bootstrapped mediation model of mPFC activity in response to childhood songs modulating negative affect through functional connectivity with the amygdala. ** p < 0.01. Error bars represent 95% confidence intervals around the mean.

### Childhood Music Regulates Negative Emotion through mPFC Circuitry

To identify how supraACC reactivity regulated negative emotion in response to childhood compared to adolescent music, functional connectivity between the supraACC region and an *a priori* bilateral anatomical amygdala seed was calculated. Stronger inverse supraACC-amygdala connectivity during childhood music relative to the adolescent music predicted decreased reported stress after childhood music (b = 0.010, t = 2.81, p = 0.011, n = 23; Figure 3C). SupraACC-amygdala connectivity mediated the association between supraACC reactivity and reported stress after childhood music (indirect effect: -0.505 (SE 0.32), CI_L95_ -1.38 – CI_U95_ -0.011, n =20; Figure 3D). That is, supraACC reactivity and SupraACC-amygdala connectivity provide a mechanism through which childhood music regulates emotion under stress.

### Early Adversity Shifts the Timing of Music Preference and Regulation Benefits

In the rodent, the timing of sensitive periods themselves has been shown to be malleable following early adversity (*18*), and recent evidence in the human suggests that early adversity may accelerate the development of the mPFC circuitry (*19*) shown here to be sensitive to childhood music. It was expected that early adversity (parental death or divorce before age 6 years old) would shift the putative mPFC sensitive period earlier in development, indexed by behavioral preference and emotion regulation benefits for the preschool music (4-6 years old at initial exposure) in the secondary sample (Figure 4A). Under stress, the adversity group showed a greater preference for preschool music relative to the comparison group (b = 0.225, t (19) = 2.25 p = 0.037, n = 20). The degree of behavioral preference for preschool music predicted subsequent increased emotion regulation only for the adversity group (group x % dwell time b = 8.065, t(20) = 2.401, p = 0.028, n = 21). That is, early adversity was associated with an earlier sensitive period in development for music’s effects, consistent with early adversity shifting developmental timing.

**Figure 4.**
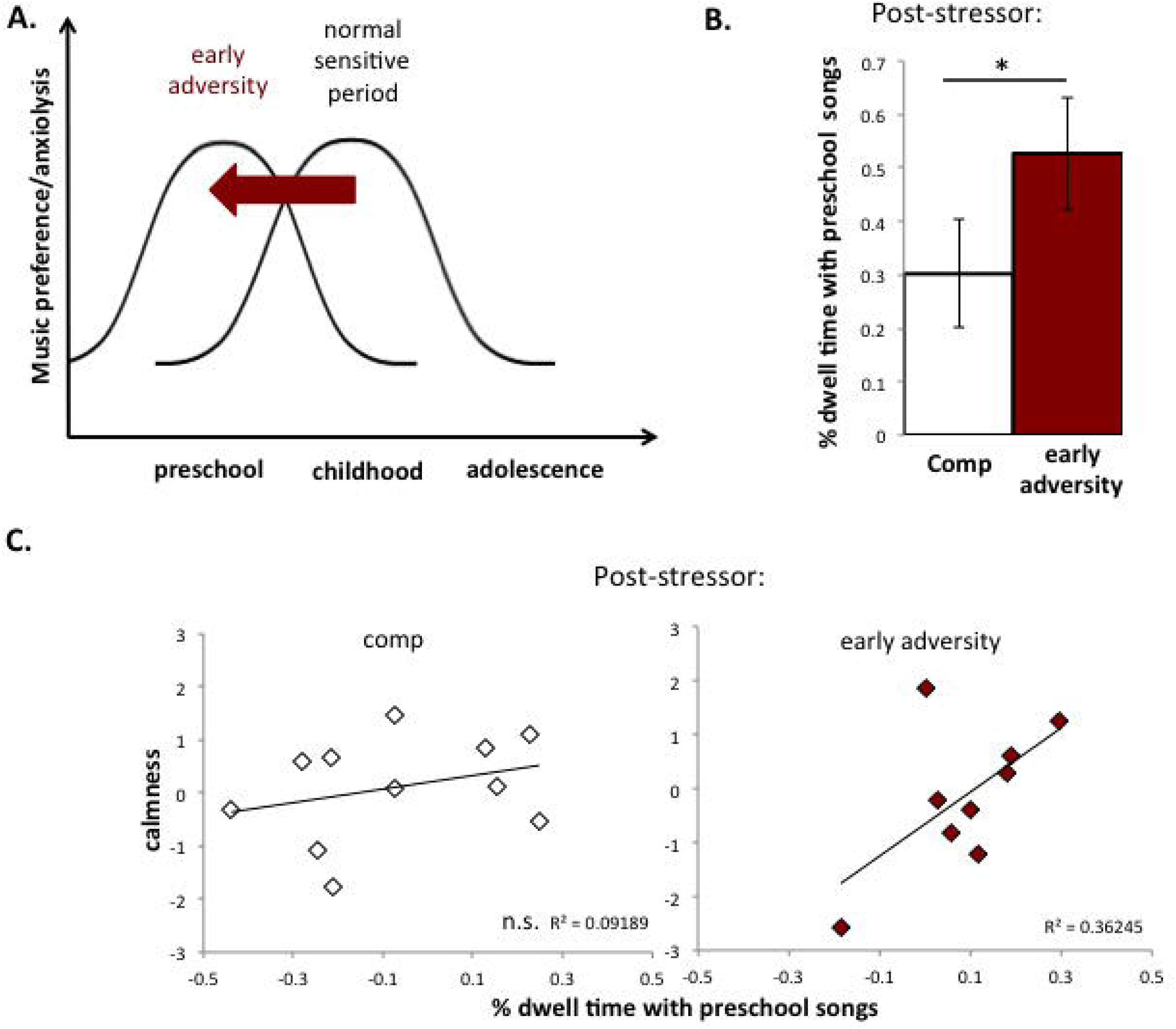
Early adversity modulates putative sensitive period for music preference and anxiolysis. A. Conceptual schematic of how early adversity is hypothesized to modulate the putative sensitive period. B. Early adversity uniquely increases preference for preschool songs relative to adolescent songs post-stressor. C. Degree of preference (% dwell time) for preschool songs post-stressor predicts reported calmness after preference assessment within the early adversity but not comparison group. Error bars represent standard error of the mean. * p < 0.05. ** p < 0.01.

An open question in neuroscience has been whether developmental sensitive periods exist for human prefrontal cortex functions. Here we tested whether childhood is a sensitive period for mPFC-mediated emotion regulation using music exposures across development. Previous work has shown that familiar music engages cortical-subcortical networks subserving affective and memory processes (*20*–*22*), and we took a developmental approach to examine music encoding. Compared to music from preschool and adolescence, we show that childhood music uniquely evokes a behavioral preference under stress and enhances activity in the mPFC, whose response and connectivity with the amygdala predict greater emotion regulation. Moreover, our findings are consistent with early adversity inducing an earlier onset of the sensitive period. The findings provide evidence that the juvenile sensitive period for music-induced regulation mediated by mPFC translates across human and rodent species (*9*).

An alternative account of the present results posits that childhood music is a regulatory signal because in comparison to adolescence, childhood is more likely to be associated with positive memories and safety, rather than constituting a sensitive period. However, adults did not report associating childhood music with significantly better life quality relative to adolescent music (t(58) = 1.434, p = 0.157, n = 59). Additionally, music from the preschool period, which should have similar positive qualities to childhood, was not preferred, except by individuals who reported early adversity. Taken together, the effect of childhood does not appear to be explained by the positive environment or safe associations of childhood.

Whether the observed sensitive period is specific to music or generalizes to other domains remains to be explored. Neither auditory cortex activity, nor auditory-mPFC functional connectivity differentiated childhood from adolescent music (Supplemental Material), suggesting the childhood-specific regulatory effects were independent of auditory cortex involvement and may generalize to other stimulus modalities (e.g. olfaction (*22*)). The current findings explain why under times of duress, individuals return to childhood nostalgia (*23*). Indeed, nostalgia, which increases supraACC activity, may be the process named to describe this sensitive period learning (*24*). The present findings reporting a developmental sensitive period for mPFC not only provide important information about the construction of the human prefrontal cortex, but also provide the opportunity and framework to identify additional sensitive periods of human associative cortex in the future.

## Methods

### Participants

Seventy young adults born in the United States of America (USA) with self-reported exposure to pop music during development (childhood (6 to 10 years old) and adolescence (15 to 19 years old) were recruited as the primary childhood-exposed group of interest. A control group of 39 young adults who had recently immigrated to the USA and reported minimal exposure to USA pop music during childhood were recruited as an additional (non-USA born) control group. All participants were between ages 18-23 years old at the time of the study and were enrolled in a Psychology course receiving credit for participation. Nine childhood-exposed group participants were excluded using a priori criteria (1 technical failure during the paradigm, 8 participants were noncompliant with the paradigm, e.g. using cell phones during the paradigm, refusing to answer any questions), therefore 61 childhood-exposed participants contributed data to analyses. Three participants were excluded from the control group using a priori criteria (2 were noncompliant with the paradigm, 1 had exposure to USA pop music during development), therefore 36 control participants contributed data to analyses. Of the included participants, 14 from the childhood-exposed group and 15 from the control group participated in an fMRI session. For the MRI visit, all participants had normal or corrected-to-normal vision. Participants were excluded from completing the MRI visit if they reported metal implants or any other contraindications to MRI.

A secondary sample of 23 participants within the same age range as the primary sample was recruited to complete a related paradigm that required exposure to USA pop music during the preschool years (4-6 years of age) in addition to the childhood and adolescent periods. Twelve participants made up an early adversity group, and had experienced either the death (n = 3) or divorce (n = 9) of their primary caregivers before age 6 years. Two participants from the early adversity group were subsequently excluded for a priori criteria (1 participant experienced their early adversity after the developmental period corresponding to the preschool-age music stimuli, 1 participant reported hating pop music and did not want to listen to it). Eleven participants made up the comparison group, and they did not experience either death or divorce of their caregivers before the age of 6 years. In total, 11 comparison participants (1 male, 10 female), and 10 early adversity participants (2 male, 8 female) contributed data to the study.

This study was approved by the Institutional Review Boards of the University of California, Los Angeles and the state of California. All participants provided informed consent for this study.

Participants came to the laboratory for up to two sessions. All participants completed the first session, where behavioral and physiological measures were collected and participants completed a stressor paradigm interleaved with music clips from childhood and adolescence. A subsample of participants from the primary sample returned for a second functional magnetic resonance imaging (fMRI) session, where they completed a related stressor paradigm administered in the scanner.

### Music Exposure Assessment

For the primary sample, all participants completed a questionnaire to assess their general exposure to USA pop music during childhood and adolescence and their associations with the music (2 participants did not report their adolescent music exposure or associations on the questionnaire). Additionally, following the stressor paradigm, participants were played 15 song clips used in the paradigm from each period (childhood and adolescence) and were asked to rate how familiar they were with each of the 30 stimuli on a 1-7 point scale (1: “never heard it before”, 7: “I can sing all the words”). The average familiarity rating for the childhood songs and adolescent songs was calculated for each participant as a measure of exposure to the stimuli specifically used in the study. The study-specific measure of childhood and adolescent stimuli exposure was significantly correlated with reported general childhood (Pearson’s r = 0.518, p = 0.000019, childhood-exposed group n = 61) and adolescent exposure (Pearson’s r = 0.340, p = 0.001, n = 95). However, a subsample of childhood-exposed participants reported high general exposure but low study specific exposure due to restricted listening to pop stations that did not play mainstream pop music (e.g. RadioDisney). Therefore, the study-specific measure of music exposure was used throughout analyses to best represent participants’ experience with the specific stimuli set. As expected, the childhood-exposed group reported significantly higher study specific childhood music exposure than the control group (t = -9.467, p = 2.8 x 10^-15^, n = 97) (Supplemental Figure 1). The childhood-exposed group also reported significantly higher study specific adolescent music exposure than the control group (t = 4.34, p = 0.000062, n = 97; Supplemental Figure 1). Within each group, participants reported significantly higher exposure to the adolescent music than to the childhood music (childhood-exposed group: t = -14.027, p = 1.335 x 10^-20^, n = 61; control group: t = -15.714, p = 1.99 x 10^-17^, n = 36, Supplemental Figure 1).

For the secondary sample, all participants completed a similar questionnaire to assess their exposure to USA pop music generally during the preschool period (ages 4-6 years old), childhood, and adolescence as well as their associations with the music from each of these periods. Following the stressor paradigm, participants were played 10 song clips from the paradigm from each period and were asked to rate how familiar they were with each of the 30 stimuli on a 1-7 point scale as in the primary sample. The average familiarity rating for the preschool, childhood, and adolescent songs was calculated for each participant as a measure of study-specific exposure to the music. The early adversity group did not significantly differ from the comparison group in either their reported exposure levels to the preschool-age music (t(20) = -0.321, p = 0.75, n = 21), or adolescent music (t(20) = -0.510, p = 0.616, n = 21). There were no group differences in reported explicit liking of preschool-age music as adults (t(19) = -0.338, p = 0.739, n = 20; 1 participant did not complete this question).

### Stimuli

Pop genre songs were selected for the study stimuli due to the transient popularity of specific songs in the genre, and therefore temporally-limited radio playtime and compact disk sales (preschool and childhood songs were released before personal devices that supported playing older songs intermixed with new songs [e.g. digital MP3 players, smartphones] were widely available). The Billboard Hot 100 chart uses sales and radio play frequency to determine song popularity, providing a metric for the timing of each song’s peak public consumption. For the purposes of this study, songs were excluded if they achieved significant global popularity, especially with non-English speaking countries (where participants in the control group largely resided). For the primary sample, thirty-six songs from 2001 were selected to be the childhood music in the study and 36 songs spanning 2010 to 2012 were selected to be the adolescent music (all songs were purchased through the iTunes music store). Songs were selected to match pop subgenres between the childhood and adolescent music sets, and the sets did not significantly differ in the proportion of major relative to minor key signatures (p = 0.562, n = 72), tempo (beats per minute) (p = 0.522), the proportion of female relative to male vocalists (p = 0.622), the proportion of solo relative to group vocals for song verses (p = 0.0626), the number of weeks songs spent on the Billboard Hot 100 chart (p = 0.140), or the peak position songs achieved on the Hot 100 chart (p = 0.839). There were significantly more group vocals for song refrains in the childhood song set than the adolescent set (p = 0.029). A 25-second clip was generated for each song using Audacity software (http://www.audacityteam.org, version 2.0.3.0) to be presented in the behavioral paradigm, and 16-second clips were generated for the fMRI paradigm. For the fMRI song clips, as they were presented without participant control, a 3-second fade-in was edited into each song clip to avoid startling participants within the scanner.

For the secondary sample, twenty-four songs from 1998 were selected to be the preschool music, and twenty-four songs each from the set of childhood and adolescent music in the primary sample were selected. Songs were selected to match pop subgenres between the preschool, childhood, and adolescent sets, and the preschool and adolescent songs did not significantly differ in the proportion of major relative to minor key signatures, tempo (beats per minute), the proportion of female relative to male vocalists, the proportion of solo relative to group vocals for song verses, or the number of weeks songs spent on the Billboard Hot 100 chart (all p > 0.10).

### Behavioral Paradigm

The stressor/song paradigm was presented to all participants using E-Prime (version 2.0) software on a computer in a quiet room, and music was played through headphones that each participant wore for the duration of the task. For the primary sample, childhood and adolescent song clips were presented to participants through two labeled “radio stations”, one of which played only childhood song stimuli (when participants were ages 6-10) and the other of which played only adolescent song stimuli (when participants were 15-19 years old) throughout the stressor paradigm. The station identity playing childhood songs was counterbalanced across participants. Participants were told that both stations played pop music. Both radio stations were presented on a choice screen, and participants could select either radio station through keypad response. Participants had to make a station choice after every song clip (i.e. they had to make a response whether they wanted to stay on the same station or switch stations). Participants were also free to change their choice at any time during a song clip through keypad response. Participants were not given any information about what specific song would play either before they selected a radio station or for the duration of the song’s playtime on the selected station. Each station cycled through the corresponding song list with random selection without replacement. If all song clips had been selected once, the station began another block of random selection without replacement.

#### Baseline song preference assessment

Participants were first presented with each radio station sequentially in the stressor paradigm. For each station, six 10-second song clips from that station’s song list were played to familiarize the participant with the station. Participants were then presented with the station choice screen and asked to choose music to listen to for five minutes. Following the preference assessment, participants were asked to report how calm they felt using a 7-point scale (1 not calm at all; 7 very calm).

#### Stressor

Participants then underwent the first block of a modified trier social stress test, consisting of math problems to solve while the experimenter watched over their shoulder and took notes on their performance (each block lasted approximately 2.75 minutes). All problems were modeled after secondary school standardized test problems, and problems had restricted timing so that it was challenging to answer many problems within the allotted time. Points were deducted if participants did not answer a problem in time before the next problem appeared, or answered a problem incorrectly, and each instance of point deduction was accompanied by an aversive buzzer sound. Participants were told their test scores would be compared with peers’ performance. After each math block, they were again asked to self-report on how calm they felt, and they were then given a five-minute “rest” period during which their music preferences were assessed. During this “rest” period, participants could freely choose to listen to either childhood or adolescent song “radio stations”. Participants reported how calm they felt again after the music preference assessment. The sequence of math stressor, reporting, music preference assessment, reporting was repeated for a total of three blocks. Participants reported feeling significantly less calm after each math stressor relative to the baseline and “rest” periods, indicating the stressor successfully modulated their affective state. After the final preference test, participants were given a surprise memory test for song clips to ensure they had been paying attention to the music during the study. For a random sample of 15 songs from each song period, they were asked to state whether or not they had heard the song during the study. Study groups did not differ in their rates of accurate recall measured through the d’ statistic (p > 0.10). The paradigm timeline is summarized in Figure 1.

The secondary sample of participants were administered the same stressor/song paradigm in a slightly longer format to accommodate the additional songs from the preschool period. Preschool, childhood, and adolescent songs were presented to participants through two pairings of radio stations, one pairing per block of the stressor paradigm. In the first pairing, participants chose between radio stations playing preschool music or adolescent music, and in the second pairing, participants chose between stations playing childhood music or adolescent music. The pairings alternated throughout the paradigm, and participants encountered each pairing twice. Station identity and the order of the pairings were counterbalanced across participants. Preference assessments were reduced to 3.75 minutes each due to time constraints. After the final preference assessment, participants were given a random sample of 10 songs from each music period during the surprise memory test.

### Depression and Anxiety Questionnaires

In addition to the music exposure questionnaire and report of early adversity, participants were asked to complete the Beck Depression Inventory-II (BDI-II), a widely used psychometric test for depression severity consisting of 21 questions (the study included 20 questions-the question about suicidality was omitted). Participants were also asked to complete the Spielberger State Trait Anxiety Inventory (STAI) for Adults, a 40 item questionnaire to evaluate state and trait anxiety levels. Two participants in the secondary sample completed neither the BDI-II nor the STAI. The BDI-II and STAI were scored in SPSS software (version 23). For the purposes of this study, the STAI trait anxiety average score was used as the measure of chronic anxiety.

### Galvanic skin conductance signal acquisition: Behavioral session

Tonic galvanic skin conductance signal was collected using two disposable isotonic electrodes attached to the inner arch and the sole of the hallux on participants’ right foot (to leave their hands free to perform tasks) prior to starting the stressor task. The galvanic skin conductance signal was recorded with 1 KHz sampling rate and amplified using a Biopac MP150 system and AcqKnowledge 4.1 software (Biopac, Goleta, CA).

### Galvanic skin conductance signal processing

Galvanic skin conductance signal was successfully acquired for 52 of the 61 childhood-exposed group participants. For each participant, galvanic skin conductance signal was processed offline with Acknowledge Software (version 4.1). The signal was filtered with a digital low-pass 1Hz filter with Blackman windowing optimized for the 2000Hz sampling rate and 1Hz cutoff and smoothed over 200 samples to remove physiological and motion-related artifact (informal communication with Biopac Systems). The filtered, smoothed signal was then manually inspected for remaining artifact and any contaminated stimulus response segments as well as any segments manually flagged for artifact during signal acquisition were removed from further analysis. Using the processed conductance signal, the signal’s slope (peak-to-trough slope) was calculated for the duration of each stressor block and for each song during the post-stress preference assessments. As participants were not given advance information about what specific song would play when they selected a radio station, slope values for songs with fewer than 6 seconds of listening time were discarded to isolate only songs that participants let play beyond the first few seconds of song recognition and/or evaluation (all 52 participants met this criteria). All participants with a minimum of 3 trials each for childhood and adolescent songs were included in analyses to ensure sufficient sampling of GSC slope values for each participant (51 of 52 participants met this criteria). The average peak-to-trough slope for all remaining songs during each post-stressor preference assessment was calculated for childhood and adolescent songs separately and averaged across post-stressor preference assessments, resulting in one slope value each for the childhood song and adolescence song conditions for each participant.

### fMRI paradigm

The songs used in the fMRI paradigm were unique to each participant, dependent on their individual song exposures. In addition to matching the childhood and adolescent songs on the behavioral session song parameters, childhood-exposed participants’ (i.e., USA born) familiarity ratings of specific songs were used to generate song sets where childhood and adolescent songs were matched for previous exposure. For the control group (i.e., non-USA born) participants, adolescent stimuli were selected to fall within the same range of previous exposure as the childhood-exposed group, and any childhood songs the participant reported familiarity with from childhood were excluded from the song set.

Participants were presented with songs through MRI-safe noise-reducing headphones in the scanner. Participants registered all responses using a 4-button MRI-compatible response box. Participants first underwent a baseline run during which childhood and adolescent songs were presented (4 clips for each condition). Following each song, participants were given a 2 second window to rate with a response box how much negative arousal they felt using a 4-point thermometer scale (from calm to a lot of negative arousal). Participants then underwent two runs of an fMRI stressor task interleaved with the songs from childhood and adolescence (as in the behavioral session). The stressor paradigm consisted of a 46-second block of timed mental math problems, followed by a 16-second block during which either a childhood or adolescent song was presented accompanied by a white fixation cross on a black screen. Participants were asked to rate negative arousal using the thermometer scale in 2-second windows following the math block and the music block (i.e. math, rate, music, rate). This sequence was repeated for a total of 5 paired math-music blocks per run (10 blocks total). During the mental math blocks, participants were given a series of problems that were timed to move from one to the next quickly with unspecified time limits for solving each problem, as before. Participants were also presented with two progress bars at the bottom of the screen, one of which they were told tracked a measure of their own speed and accuracy, the other of which represented the average performance for a peer group. As an additional stressor, the bars were presented so that the participant often appeared to be performing more poorly than the peer group average. The fMRI paradigm is summarized in Figure 1.

### fMRI acquisition

All participants were scanned with a Siemens Trio 3.0-Tesla MRI scanner using a standard 12-channel radiofrequency head coil. Two functional scans of T2*-weighted echoplanar images (interleaved) were collected at an oblique angle of ∼15° to 30° (selected per participant to minimize signal drop-out for their scans) (186 volumes/run; TR, 2000 ms; TE, 30 ms; flip angle, 75°; matrix size, 64 × 64; field of view (FOV), 192 mm; 34 slices; 4 mm slice thickness; skip 0 mm; 1420Hz bandwidth; 5 song blocks per run alternating between song periods throughout, condition order counterbalanced across participants in each group separately). A whole brain, high resolution, T1-weighted anatomical scan (MP-RAGE; 256 × 256 in-plane resolution, 250 mm FOV; 176 mm × 1 mm sagittal slices oversampled by 18.2%; 200Hz bandwidth) was acquired for each participant for registration and localization of functional data to Talairach space (*1*).

### fMRI Data processing

The functional imaging data were preprocessed and analyzed with the Analysis of Functional NeuroImages (AFNI) software package (*2*). For each participant’s images, preprocessing included discarding the first 4 functional volumes to allow for BOLD signal stabilization, correction for slice acquisition dependent time shifts per volume, rigid body translation and rotation from each volume to the first volume to generate 6 within-subject regressors, generating registration parameters to the participant’s anatomical scan, and spatial smoothing. Data were smoothed to an 8mm isotropic full-width half maximum smoothness using 3dBlurToFWHM (i.e., various smoothing kernels were used across participants to achieve the same effective spatial smoothness of 8mm) to reduce differences in smoothness across participants due to different levels of motion artifact (Scheinhost et al 2014). To allow for comparisons across individuals, timecourses were then normalized to percent signal change, functional data were registered to the anatomical scan using the parameters prior to smoothing, and the anatomical and functional scans were transformed to the standard coordinate space of Talairach and Tournoux (*1*) with align_epi_anat.py. Transformations on the functional scans (to the participant’s anatomical scan and to the standard Talairach and Tournoux template space) were combined into a single transformation within align_epi_anat.py to minimize the amount of interpolation applied to the functional data. Talairach-transformed images had a resampled resolution of 3mm^3^. Functional runs were concatenated before creating individual-level models for each participant for stimulus-elicited activity and connectivity analyses.

### Motion Corrections

Consistent with recent recommendations, a strict motion-censoring limit was applied so that any timepoint and the immediately preceding timepoint were both censored if the Euclidean norm of the scan-to-scan motion parameters across the 6 rigid-body parameters exceeded 0.25 mm/degrees (Siegel et al., 2014; Power et al. 2015). One participant in the childhood-exposed group was excluded from further analysis for excessive motion (participant’s Euclidean norm of scan-to-scan motion before censoring >0.50mm/degrees). At the within-subject level of analysis, 6 rigid-body motion regressors and the 6 backwards temporal derivatives of those regressors were included in all regressions to additionally correct for head motion artifacts (*3*, *4*).

### Stimulus-Elicited Reactivity Analysis

A GLM analysis was performed in AFNI for each participant using 3dDeconvolve to assess stimulus-elicited activity changes across the whole brain. In addition to regressors for each stimulus type (stressor task, self-report of emotion, childhood song, adolescence song), timecourses for 12 motion regressors (6 rigid-body regressors and their 6 backwards temporal derivatives) and timecourses from eroded ventricle, eroded white matter masks, and whole-brain (global signal) masks were included as physiological nuisance covariates. The GLM analyses fit the percentage signal change time series to each regressor, and linear, quadratic, and cubic trends were modeled for the time series of each voxel to control for correlated drift.

### Stimulus-Elicited Connectivity Analysis

Generalized psychophysiological interaction (gPPI) analysis was conducted to assess stimulus-dependent medial prefrontal cortex (mPFC) connectivity changes across the whole brain in each participant (Friston et al., 1997). Main effects of the psychological (stimulus) regressors and physiological (the functionally-defined mPFC stimulus-elicited activity timecourse) regressor were controlled for through inclusion in this gPPI analysis. Four psychological (stimulus) regressors modeled whether a given block consisted of the stressor task, self-report of emotion, childhood song, or adolescence song. The physiological (seed region time series) regressor was the time series for the mPFC seed region after regressing out fixation and drift (by modeling linear and quadratic trends for the timeseries). Four interaction regressors modeled the interaction of the psychological regressors and the physiological regressor, such that each interaction regressor identified regions whose time series correlated in a stimulus-dependent manner with the mPFC time series. The gPPI GLM analysis was performed in AFNI using 3dDeconvolve for each participant with regressors for stimuli, mPFC seed region timecourse, interactions of each stimulus and seed timecourse, timecourses for eroded ventricle and eroded white matter masks and whole-brain masks as physiological nuisance covariates, and 12 motion regressors (6 rigid-body regressors and their 6 backwards temporal derivatives). The GLM analyses fit the percentage signal change time series to each regressor, and linear, quadratic, and cubic trends were modeled for the time series of each voxel to control for correlated drift. A general linear test was calculated within the GLM for a second-order contrast of mPFC x childhood stimuli relative to mPFC x adolescent stimuli conditions to compare how mPFC connectivity modulated by childhood songs differed from mPFC connectivity modulated by adolescent songs.

### Statistical Analyses

All statistical analyses were run in SPSS software (version 23). A priori outliers were determined to be any data point with a studentized deleted residual value in the analysis of interest greater than 3.

### Behavioral Analyses

For the primary sample, for all preference assessments in the stressor paradigm, the percent of time the participant spent listening to songs on the childhood station relative to the adolescent station was used as the measure of preference for childhood songs in the study. The preference score for childhood songs was averaged across the three post-stressor preference assessments. The childhood-exposed group included participants with a wide range of childhood song exposures, and post hoc analyses tested how varying exposure within this group related to behavioral preferences compared to the control group; the child-exposed group was median-split on the exposure scale into a low-exposure subgroup (exposure scores 1-3.49, n = 22) and a higher-exposure subgroup (scores 3.5 -7, n = 39). Repeated measures ANCOVA was conducted on the percent of time spent with the childhood songs in the pre-stressor baseline and post-stressor assessment periods (within-participant factor) across the control, low-exposure subgroup, and higher-exposure subgroup of the exposed group (between-participant factor). The ANCOVA controlled for individual differences in exposure to the adolescent songs (between-participant covariate). Post-hoc simple effect tests used the Bonferroni correction for multiple comparisons so the family-wise error rate was less than 0.05.

To examine group differences in reported emotion regulation following the preference assessments, the change in reported calmness was first calculated (average report of post-preference assessments – average report of post-stressor assessments, excluding the baseline post-preference assessment score). Linear regression was performed with group (childhood-exposed group or control group) predicting the calmness change score, with each participant’s adolescent music exposure score and post-stressor assessment report included as covariates.

Nonparametric empirical bootstrapping mediation (10,000 iterations) with bias-corrected confidence intervals was performed using the PROCESS macro in SPSS (version 23) as a robust approach for smaller sample sizes to test the mediated (indirect) effect of the behavioral preference for childhood songs post-stressor on the relation between group (childhood-exposed or control) and the calmness change score, with adolescent song exposure and post-stressor assessment report included as covariates as before (*7*). A second mediation was performed with PROCESS within the childhood-exposed group participants only, testing the mediated effect of behavioral preference for childhood songs post-stressor on the relation between childhood song exposure score (continuous measure), and the calmness change score, controlling for the same covariates as before.

For the secondary sample, the early adversity group had significantly higher self-reported levels of depression on the BDI-II (t (19) = -2.86, p = 0.010, n = 20) and trait anxiety on the STAI (t(19) = -2.84, p = 0.01, n = 20) than the comparison group. To ensure the analyses examining music preference were not due to these differences in concurrent psychopathology, the grouping variable and the preschool music preference variable were each residualized on the trait anxiety and depression scores.. To test whether the early adversity group differed from the comparison group in their preference for the preschool music, the residuals for the group and preference variables were then fed into a Pearson’s correlation test. To examine group differences in reported emotion regulation following the music preference assessments, a linear regression was performed with group (early adversity or comparison), preschool music preference score, and the interaction between group and preschool music preference score predicting self-reported calmness scores. One early adversity participant was removed as an overly-influential outlier using a priori criteria (participant’s residual > 4). Post-hoc simple slope tests were conducted for each group separately.

### Physiological Statistical Analyses

Repeated measures ANCOVA was conducted with childhood and adolescent exposure scores (between-participant continuous measure factors) predicting the mean post-stressor galvanic skin conductance (GSC) peak-to-trough slope for childhood songs and adolescent songs (within-participant factors), controlling for participant differences in GSC slope during the stressor (between-participant covariate). Post-hoc simple effect tests used the Bonferroni correction for multiple comparisons so the family-wise error rate was less than 0.05. Simple effects of GSC slopes for childhood and adolescent songs were evaluated at childhood and adolescent exposure scores equal to 2 (low exposure), 4 (medium exposure; the scales’ median value), and 6 (high exposure), scores that were within the range of the group’s actual exposure scores.

Post-stressor GSC slopes for childhood and adolescent songs were then entered into linear regression with the post-stressor assessment calmness report (covariate of no interest) to predict the calmness change score. As the adolescent GSC slope was not significantly related to the calmness change score in that model, it was removed and the linear regression was re-run.

### fMRI Reactivity Statistical Analyses

Individual-level regression coefficients for reactivity differences post-stressor during childhood compared with adolescent songs (Childhood Songs – Adolescent Songs [CS – AS] contrast) were submitted to the group level and tested for group differences in the CS – AS contrast (childhood-exposed group CS – AS – control group CS – AS), controlling for differences in participants’ adolescent song exposure scores in a whole-brain voxelwise ANCOVA with AFNI’s 3dTtest++ program. Uncorrected voxel significance thresholding was set to p < 0.005. AFNI’s 3dFWHMx and 3dClustSim programs were used to correct for multiple comparisons to achieve a family-wise error rate of alpha < 0.01 (version compiled after the correction of a long-standing bug in 3dClustSim in 2015). AFNI’s 3dFWHMx program was run for each participant to estimate the individual-level regression’s residual smoothness in the x, y, and z directions, and participant estimates were averaged together in each direction (mean smoothness across participants in each direction: x = 7.39mm, y = 7.37mm, z = 7.45mm). The residual smoothness estimates were then submitted to AFNI’s 3dClustSim for 10,000 Monte Carlo simulations. The critical cluster threshold to achieve a family-wise error rate of alpha < 0.01 was set to 45 voxels as determined by the 3dClustSim simulations. Parameter estimates for clusters surviving thresholding were extracted for the childhood song (relative to baseline, CS - baseline) condition and the adolescent song (relative to baseline, AS - baseline) condition and residualized for differences in adolescent song exposure to match the model run in AFNI. Post-hoc Student’s T tests within each group separately were conducted on each condition’s residualized reactivity (relative to baseline) and compared to 0. Post-hoc repeated measures ANCOVA were conducted within each group separately on CS - baseline and AS –baseline reactivity (within-participant factor), controlling for adolescent exposure scores (between-participant covariate) to test whether conditions significantly differed in reactivity within each group. Linear regressions (effectively correlations) with CS – baseline reactivity predicting reports of childhood song exposure in adolescence or current liking for childhood songs were conducted to test potential confounding effects.

Reactivity parameter estimates for the CS – baseline and AS – baseline conditions were also extracted for several anatomically-defined a priori regions of interest based on prior studies of music-evoked fMRI activity (*8*–*11*). Specifically, reactivity estimates were extracted for the bilateral amygdala (defined with the Talairach Daemon atlas included in AFNI) (*8*–*10*, *12*, *13*), bilateral hippocampus (defined with the Talairach Daemon atlas included in AFNI)(*9*, *12*, *13*), bilateral ventral striatum (defined with the Oxford-GSK-Imanova structural striatal atlas, (*8*, *9*, *13*, *14*) included in FSL (FMRIB, Oxford, UK), and bilateral auditory cortex (defined as Heschel’s gyrus and including regions TE 1.0, TE 1.1, TE 1.2 with the Juelich histological atlas included in FSL, (*15*). Repeated measures ANCOVA was conducted for each a priori region on CS – baseline and AS – baseline reactivity (within-participant factor) as a function of group (childhood-exposed or control, between-participant factor), controlling for adolescent exposure scores (between-participant covariate).

To examine how the whole-brain thresholded regions differentially related to behavior, repeated measures ANCOVA and post-hoc simple effects’ tests were conducted on the regions’ reactivity in the CS – baseline condition (within-participant factor) and reported negative arousal following post-stressor childhood songs (between-participant continuous factor), controlling for negative arousal reported following pre-stressor childhood songs (baseline arousal level). To examine how the whole-brain thresholded regions differentially related to participants’ preference for one condition relative to the other (indexed by their choice behavior during the behavioral session), repeated measures ANCOVA and post-hoc simple effects’ tests were also conducted on the regions’ reactivity for the CS –AS contrast (within-participant factor) and percent time spent with childhood songs relative to adolescent songs post-stressor during the behavioral stressor task (between-participant continuous factor).

### fMRI Connectivity Statistical Analyses

Individual-level gPPI regression coefficients for the second-order contrast of mPFC x childhood songs compared to mPFC x adolescent songs (mPFCxCS – mPFCxAS contrast) were submitted to a group-level whole-brain voxelwise ANCOVA with AFNI’s 3dTtest++ program testing differences between groups (childhood-exposed group [mPFCxCS – mPFCxAS] – control group [mPFCxCS – mPFCxAS]) and controlling for differences in participants’ adolescent music exposure scores. Uncorrected voxel significance thresholding was set to p < 0.02 (a more lenient threshold was used given gPPI’s higher rate of false negatives, O’Reilly et al., 2012). AFNI’s 3dFWHMx and 3dClustSim programs were used to correct for multiple comparisons to achieve a family-wise error rate of alpha < 0.05. The mean residual smoothness across participants in each direction for the supragenual-anterior cingulate seeded gPPI was x = 7.4mm, y = 7.38mm, z = 7.44mm, yielding a critical cluster threshold of 75 voxels as determined by the 3dClustSim simulations. The mean residual smoothness across participants in each direction for the subgenual-anterior cingulate seeded gPPI was x = 7.4mm, y = 7.4mm, z = 7.45mm, yielding a critical cluster threshold of 76 voxels as determined by the 3dClustSim simulations. Parameter estimates for clusters surviving thresholding were extracted for the mPFCxCS condition and the mPFCxAS condition and residualized for differences in adolescent music exposure (residuals from regression of adolescent exposure score on parameter estimates). Post-hoc Student’s T tests within each group separately were conducted on each condition’s residualized connectivity (relative to baseline) and compared to 0. Post-hoc repeated measures ANCOVA were conducted within each group separately on the connectivity conditions (within-participant factor), controlling for adolescent exposure scores (between-participant covariate) to test whether conditions significantly differed in connectivity within each group. Linear regressions tested associations between connectivity conditions and reported post-song negative arousal or reported positive associations with the stimuli during development.

Connectivity parameter estimates for the mPFCxCS condition and mPFCxAS condition were also extracted for the anatomically-defined a priori amygdala, ventral striatum, hippocampus, and auditory cortex regions of interest. Repeated measures ANCOVA was conducted for each a priori region on the connectivity conditions (within-participant factor) as a function of group (between-participant factor), controlling for adolescent exposure scores (between-participant covariate). Linear regressions tested associations between the mPFCxCS – mPFCxAS contrast parameter estimates and reported post-song negative arousal, controlling for pre-stressor reported negative arousal (baseline arousal level). Nonparametric empirical bootstrapping mediation (10,000 iterations) with bias-corrected confidence intervals was performed to test the mediated (indirect) effect of mPFC-amygdala connectivity in the childhood song condition relative to the adolescent song condition (CS – AS) on the association between mPFC reactivity in the childhood song condition relative to the adolescent song condition (CS – AS) and reported negative arousal following post-stressor childhood songs, controlling for negative arousal reported pre-stressor (baseline arousal level).

## Supplemental Results

### Physiological and behavioral regulation results

We further tested whether the degree of physiological and behavioral regulation during the stressor paradigm were related. The degree of physiological regulation during childhood songs was significantly associated with self-reported emotion regulation, as more negative GSC slope during childhood songs was related to subsequent increases in reported calmness (b = -39.72, t = -2.224, p = 0.031, n = 50), controlling for differences in individuals’ reported calmness pre-preference assessment. The association between GSC slope during childhood music and the change in calmness remained significant (b= - 38.95, t = -2.168, p = 0.035, n = 50) after including GSC slope during adolescent songs in the model (GSC slope during adolescent songs did not significantly predict change in calmness, b= -11.02, t = -0.784, p = 0.437, n = 50).

### FMRI reactivity results

#### SubACC Region: Relative Preference/Hedonic Value

Under conditions of stress, the childhood-exposed and control groups showed an additional significant difference in fMRI signal reactivity to childhood relative to late adolescent music in a subgenual anterior cingulate cortex region (subACC) (center of mass: x = 3, y = 36, z = 0, cluster extent = 47 voxels) (voxelwise uncorrected p < 0.005, FWE alpha < 0.01, critical cluster size 45 voxels; Supplemental Figure 2). In contrast to the supraACC region presented in the primary results, post-hoc tests on subACC reactivity within each group separately did not identify any stimuli conditions significantly different from 0 (childhood-exposed group: CS – baseline condition p = 0.065, T = 2.029, n= 13; all other conditions p > 0.1). Moreover, subACC reactivity did not differ between childhood and adolescent songs in either group (childhood-exposed group (F (1, 11) = 1.299, p = .279, n =13; control group (F(1, 13) = 0.313, p = 0.585, n = 15). That is, no specific condition or group’s reactivity contributed to the observed group difference in the subACC region beyond the relative differences between conditions.

To test whether mPFC activity was associated with the emotion regulating effects of childhood music, associations between emotion regulation and the reactivity of the two ACC regions (i.e., supraACC and subACC) whose activity differentiated childhood from adolescent music were examined. There was a significant dissociation between the two ACC regions’ reactivity and negative affect following childhood music (emotion regulation x region interaction: F (1, 18) = 4.95, p = 0.039, n = 21). As reported in the primary results, greater supraACC reactivity predicted decreased negative arousal post-stressor (negative arousal b: -0.249, t = -2.352, p = 0.030), consistent with the supraACC region facilitating the childhood music’s regulatory effects. In contrast, subACC reactivity was not significantly associated with negative arousal ratings (b: -0.088, t = - 0.826, p = 0.420). Instead, a significant interaction between ACC region reactivity (CS – AS contrast) and post-stressor choice behavior (percent time spent with childhood relative to adolescent songs during the behavioral session) (F (1, 25) = 5.09, p = 0.033, n = 27) demonstrated that subACC, but not supraACC, reactivity was significantly positively related to the degree of participant’s preference for one music period over the other (with increased reactivity during the preferred music condition relative to the other condition) (degree of preference beta for subACC region: 0.579, t= 2.91, p = 0.008; degree of preference beta for supraACC region: 0.274, t = 1.73, p = 0.095). These results are consistent with prior reports that the subACC tracks music pleasantness and musical preference regardless of the age at music exposure, capturing a component of the neural response to music distinct from the emotion regulation effects seen for the supraACC mPFC region (*8*, *16*–*19*).

#### A priori control region reactivity

Reactivity to childhood and adolescent songs were also extracted from bilateral anatomically defined a priori regions of interest consisting of the amygdala, hippocampus, ventral striatum, and auditory cortex, and analyzed offline. Participant differences in adolescent music exposure were controlled for in all analyses.

#### Hippocampus

There was a significant group difference in hippocampal reactivity, such that the childhood-exposed group had higher hippocampal reactivity than the control group (control group mean estimate = -0.096 CI -.153 to -0.039, exposure group mean estimate = 0.024, CI -0.04 to 0.088, F (1, 24) = 7.706, p = 0.01, n = 27) regardless of music stimuli. There were no significant condition or condition x group hippocampal reactivity differences (condition: F (1, 24) = 0.135, p = 0.717, n = 27; Group x Condition: F (1, 24) = 0.033, p = 0.857, n = 27).

#### Ventral striatum

Ventral striatum reactivity did not show any significant differences by condition (F(1, 24) = 0.016, p = 0.901, n = 27), group (F(1, 24) = 1.311, p = 0.264), or Group x Condition (F(1, 24) = 4.093, p = 0.054).

#### Amygdala

Amygdala reactivity did not show any significant differences by condition (F (1, 25) = 0.508, p = 0.483, n = 28), group (F (1, 25) = 1.078, p = 0.309, n = 28), or Group x Condition (F(1, 25) = 0.074, p = 0.787, n = 28).

#### Auditory cortex

Auditory cortex reactivity did not show any significant differences by condition, (F (1, 24) = 0.021, p = 0.887, n = 27), group (F(1, 24) = 2.163, p = 0.154, n = 27), or Group x Condition (F(1, 24) = 0.269, p = 0.608, n = 27).

### FMRI functional connectivity results

#### SupraACC Connectivity

A whole-brain gPPI analysis identified regions that were differentially coupled with the supraACC region during childhood relative to adolescent songs in the childhood-exposed group compared to the control group (childhood-exposed CS – AS – control CS – AS), controlling for participant differences in adolescent music exposure. A significant group difference in supraACC connectivity was observed with a region centered in parahippocampal gyrus (and including posterior cingulate, center of mass: x = 2, y = -34, z = 7 (right parahippocampal gyrus), cluster extent = 82 voxels, voxelwise uncorrected p< 0.02, FWE alpha < 0.05, critical cluster size = 75 voxels; Supplemental Figure 3). Post-hoc tests identified that only for the childhood condition (relative to baseline), the childhood-exposed group had significantly positive supraACC-parahippocampal connectivity and the control group had significantly negative connectivity (mean childhood-exposed group CS-baseline connectivity = 19.54, CI_L95_ 0.614 – CI_U95_ 38.47, t= 2.25, p = 0.044, n = 13; mean control group CS-baseline connectivity: -16.94, CI_L95_ -28.82 – CI_U95_ -5.05, t = -3.06, p = 0.009, n = 15; mean childhood-exposed and control groups’ AS-baseline connectivity = all p > 0.2). Neither group had significantly different supraACC-parahippocampal connectivity during childhood relative to adolescent songs (all p >0.1).

#### SupraACC Connectivity and Emotion Regulation

SupraACC-parahippocampal connectivity during childhood relative to adolescent songs was not significantly related to self-report of negative affect following the post-stressor childhood songs (whether or not the model controlled for any relation with negative arousal pre-stressor) (b = -0.003, t = -1.07, p = 0.299, n = 22). However, regardless of group, increasingly positive connectivity during the adolescent songs (AS – baseline) was significantly associated with greater reported positive associations with the adolescent songs during adolescence (b: 0.018, t = 2.42, p = 0.024, n = 26; Supplemental Figure 3). The SupraACC-parahippocampal findings are consistent with prior studies showing that stimuli evoking autobiographical memories activate these regions (Svoboda et al., 2006, Gilboa et al., 2004, Levine et al., 2004, Koeschl 2010). This functional circuit was not directly related to the emotion regulation effects observed, but may represent musical context retrieval or autobiographical memory retrieval that co-occurs during song presentation and is not specific to a specific developmental period.

Associations between SupraACC connectivity with the a priori regions and negative arousal were also tested. SupraACC-hippocampus connectivity was not tested given the significant whole-brain SupraACC-parahippocampal connection already tested above. SupraACC connectivity with the anatomically defined ventral striatum, and SupraACC connectivity with the auditory cortex during childhood relative to adolescent songs was not significantly related to self-report of negative arousal following post-stressor childhood songs (regardless of whether the statistical model controlled for any relation with negative arousal pre-stressor) (SupraACC-striaum t = 0.110, p = 0.913, n = 21; SupraACC-auditory cortex t = 0.556, p = 0.578, n = 21). It should be noted that although the SupraACC-amygdala connectivity analysis reported in the main text was an a priori analysis, those findings also survive Bonferroni correction for multiple comparisons in the context of these additional connectivity analyses.

#### SubACC Connectivity

A whole-brain gPPI analysis (childhood-exposed group (CS – AS contrast) – control group (CS – AS contrast)) identified a significant group difference in subACC connectivity to childhood relative to adolescent music with a region including the dorsal striatum(DS) and insula (center of mass: x = 30, y = -11, z = 13 (right hemisphere insula/claustrum), cluster extent = 80 voxels, voxelwise uncorrected p < 0.02, FWE alpha < 0.05, critical cluster size = 76 voxels) controlling for differences in adolescent music exposure. Post-hoc tests within each group showed that subACC-DS/insula connectivity only significantly differed from 0 for the childhood-exposed group during the childhood songs (mean childhood-exposed group CS-baseline connectivity = -8.18, CI_L95_ -15.56 – CI_U95_ -0.808, t = -2.42, p = 0.032, n = 13; mean control group CS-baseline connectivity = 7.09, CI_L95_ -0.230 – CI_U95_ 14.417, t = 2.077, p = 0.057, n = 15; mean childhood-exposed and control groups’ AS-baseline connectivity = all p > 0.2). The childhood-exposed group had significantly more negative connectivity during the childhood relative to the adolescent songs (exposed group: F (1, 11) = 18.748, p = 0.001, n = 13; control group: F(1, 13) = 0.200, p = 0.662, n = 15). SubACC-DS/insula connectivity during childhood relative to adolescent songs was not significantly related to reported negative arousal following childhood songs, (whether or not the model controlled for any relation with negative arousal pre-stressor) (b = -0.0001, t = -0.042, p = 0.967, n = 21). Instead, subACC-DS/insula connectivity during childhood relative to adolescent songs was significantly related to post-stressor choice behavior (percent time spent with childhood relative to adolescent songs during the behavioral session), consistent with the subACC network reflecting relative preferences (subACC-DS/insula connectivity b coefficient: - 0.006, t = -2.44, p = 0.022, n = 28).

SubACC connectivity with each of the a priori anatomical regions during childhood relative to adolescent songs (CS – AS contrast) was not significantly related to reported negative arousal following childhood songs, (whether or not the model controlled for any relation with negative arousal pre-stressor) (subACC-amygdala b = 0.007, t = 1.70, p = 0.105, n = 22; subACC-auditory cortex b = -0.004, t = -1.121, p = 0.276, n = 22; subACC-ventral striatum b = -0.005, t = -1.47, p = 0.158, n = 22; subACC-hippocampus b = 0.002, t = 0.463, p = 0.648, n =22).

It should be noted that prior studies examining brain responses to self-relevant music, musical preference, and music familiarity have reported distributed cortical and subcortical responses beyond those found in the present study, including lateral prefrontal cortex, cerebellum, and temporal regions (e.g. (*8*, *9*). The fMRI reactivity and connectivity results observed in the present study may be more limited as the primary contrasts of interest involved differences between familiar songs across developmental periods.

---

**Figure S1.**
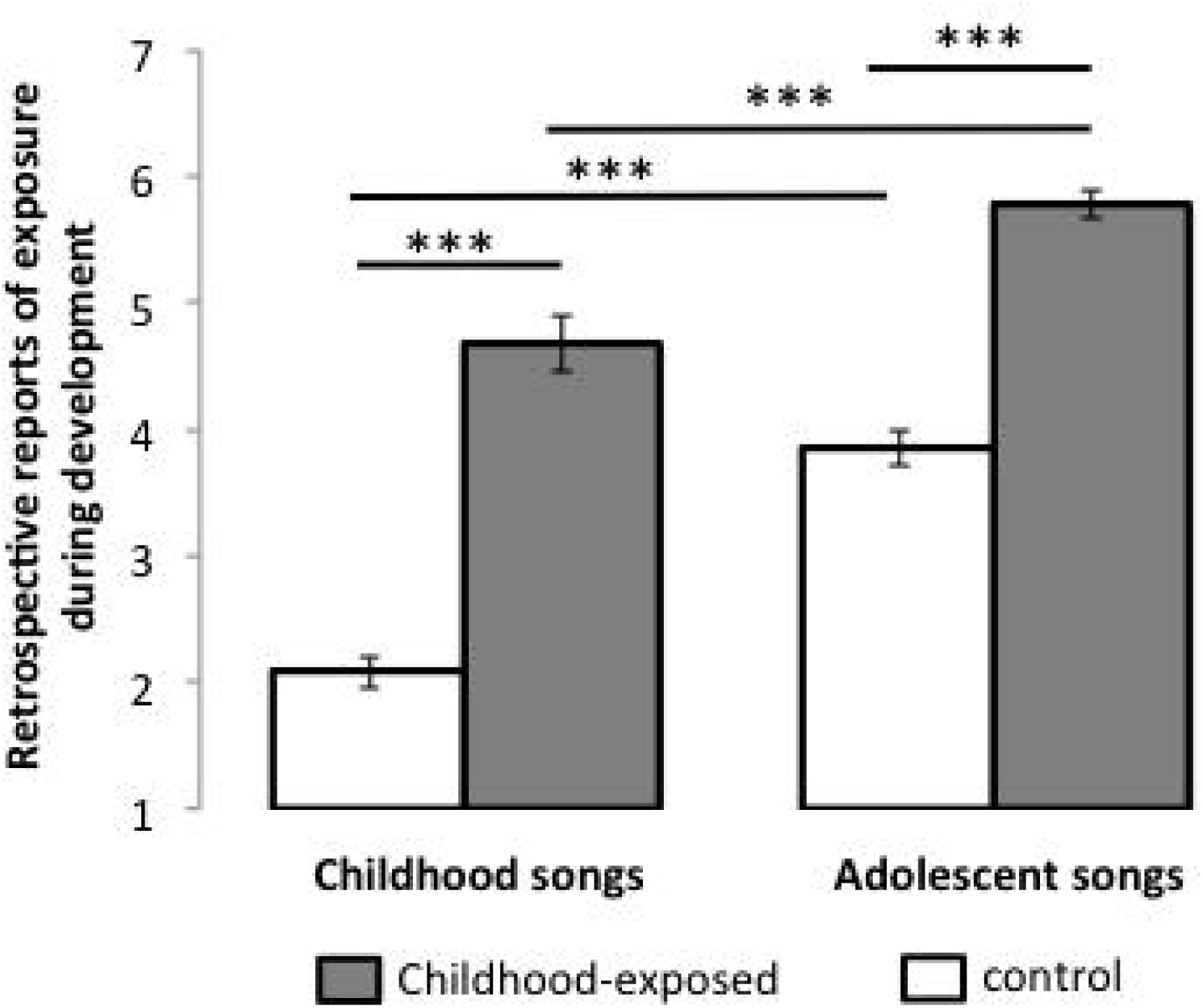
Music exposure during development. Retrospective self-report in young adulthood of exposure to the study’s pop songs during childhood (childhood songs) or adolescence (adolescent songs) in the USA-born (childhood-exposed group) and recent immigrant (control group) participants. *** p < 0.001. Error bars represent standard error of the mean.

**Figure S2.**
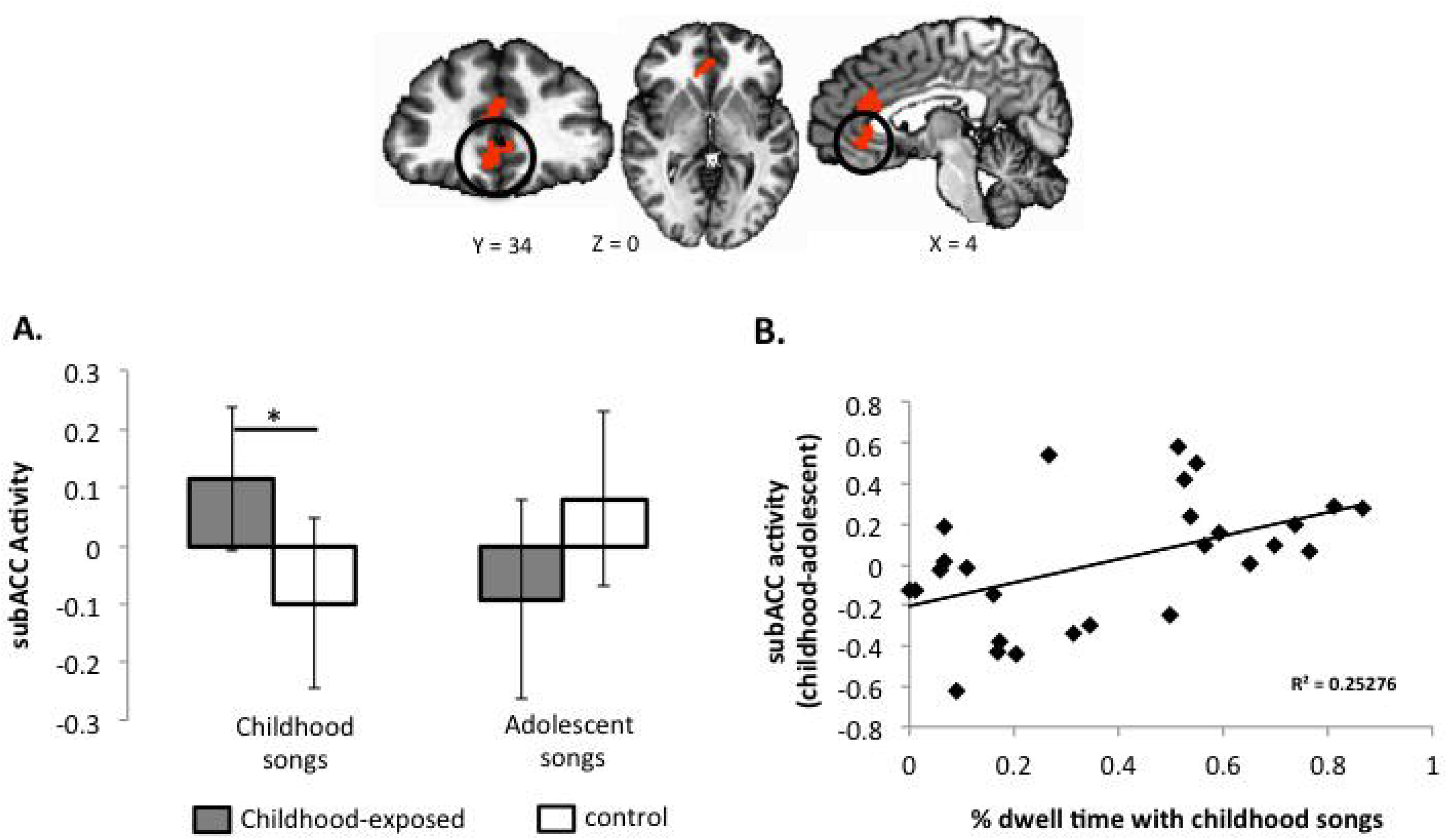
Medial pre-frontal cortex response depends on developmental exposure and is associated with behavioral preferences for childhood music. Whole-brain functional magnetic resonance imaging (fMRI) analysis showing subgenual anterior cingulate (subACC) medial prefrontal cortex region (in black circle) differentiates childhood songs between childhood-exposed and control (no childhood-exposure) groups. P < 0.005, alpha < 0.01, corrected. A. fMRI parameter estimates are shown for each condition (versus implicit baseline). B. Behavioral preference (% dwell time) for childhood relative to adolescent music (x-axis) is associated with medial prefrontal cortex reactivity (childhood versus adolescent songs; y-axis). * p < 0.05. Error bars represent 95% confidence intervals around the mean.

**Figure S3.**
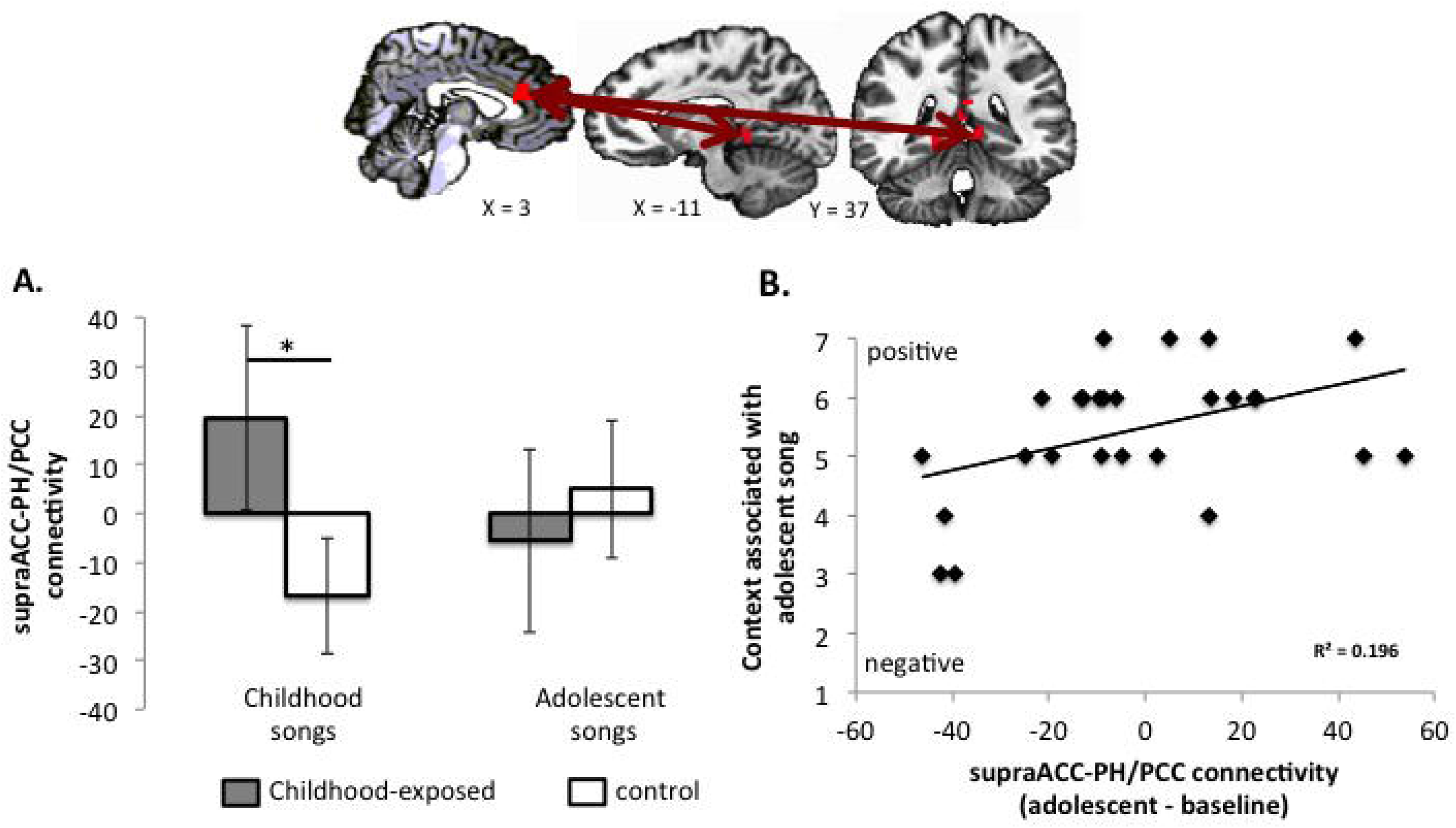
Medial pre-frontal cortex-parahippocampal circuitry responds to childhood music and is associated with recalled developmental context. Whole-brain functional magnetic resonance imaging (fMRI) analysis showing supragenual anterior cingulate (supraACC) medial prefrontal cortex – parahippocampal (PH)-posterior cingulate cortex (PCC) functional connectivity differentiates childhood songs between childhood-exposed and control (no childhood-exposure) groups. P < 0.02, alpha < 0.05, corrected. A. fMRI connectivity parameter estimates are shown for each condition (versus implicit baseline). Functional connectivity between the mPFC, PH, and PCC regions during adolescent songs (versus implicit baseline) (x-axis) is associated with the self-reported context for those songs during adolescence (y-axis). Contexts were evaluated by valence from negative (i.e. worst time of my life) through positive (i.e. best time of my life) conditions during development. * p < 0.05. Error bars represent 95% confidence intervals around the mean.

## References

1. T. K. Hensch, Critical period plasticity in local cortical circuits. Nat. Rev. Neurosci. 6, 877–888 (2005).

2. G. J. Quirk, J. S. Beer, Prefrontal involvement in the regulation of emotion: convergence of rat and human studies. Curr. Opin. Neurobiol. 16, 723–7 (2006).

3. M. R. Delgado, A. Olsson, E. A. Phelps, Extending animal models of fear conditioning to humans. Biol. Psychol. 73, 39–48 (2006).

4. M. R. Milad et al., Recall of fear extinction in humans activates the ventromedial prefrontal cortex and hippocampus in concert. Biol. Psychiatry. 62,446–54(2007).

5. M. R. Milad, G. J. Quirk, Neurons in medial prefrontal cortex signal memory for fear extinction. Nature. 420, 70–4 (2002).

6. E. A. Phelps, M. R. Delgado, K. I. Nearing, J. E. LeDoux, Extinction learning in humans: role of the amygdala and vmPFC. Neuron. 43, 897–905 (2004).

7. T. D. Wager et al., Brain mediators of cardiovascular responses to social threat: part I: Reciprocal dorsal and ventral sub-regions of the medial prefrontal cortex and heart-rate reactivity. Neuroimage. 47, 821–35 (2009).

8. C. A. Hartley, F. S. Lee, Sensitive periods in affective development: nonlinear maturation of fear learning. Neuropsychopharmacology. 40, 50–60 (2015).

9. E.-J. Yang, E. W. Lin, T. K. Hensch, Critical period for acoustic preference in mice. Proc. Natl. Acad. Sci. U. S. A. 109 Suppl, 17213–20 (2012).

10. S. S. Pattwell et al, Dynamic changes in neural circuitry during adolescence are associated with persistent attenuation of fear memories. Nat. Commun. 7, 11475 (2016).

11. M. Wu et al., Age-related changes in amygdala-frontal connectivity during emotional face processing from childhood into young adulthood. Hum. Brain Mapp. 37,1684–1695 (2016).

12. S. B. Perlman, K. A. Pelphrey, Developing connections for affective regulation: age-related changes in emotional brain connectivity. J. Exp. Child Psychol. 108, 607–20 (2011).

13. D. G. Gee et al, Maternal Buffering of Human Amygdala-Prefrontal Circuitry During Childhood but Not During Adolescence. Psychol. Sci. 25 (2014), doi:10.1177/0956797614550878.

14. L. J. Gabard-Durnam et al., Stimulus-Elicited Connectivity Influences Resting- State Connectivity Years Later in Human Development: A Prospective Study. /. Neurosci. 36, 4771–4784 (2016).

15. L. J. Gabard-Durnam et al., The development of human amygdala functional connectivity at rest from 4 to 23years: A cross-sectional study. Neuroimage. 95 (2014), doi:10.1016/j.neuroimage.2014.03.038.

16. D. G. Gee et al, A developmental shift from positive to negative connectivity in human amygdala-prefrontal circuitry. J. Neurosci. 33, 4584–93 (2013).

17. N. Tottenham, L. J. Gabard-Durnam, The developing amygdala: a student of the world and a teacher of the cortex. Curr. Opin. Psychol. 17, 55–60 (2017).

18. B. L. Callaghan, R. Richardson, Maternal separation results in early emergence of adult-like fear and extinction learning in infant rats. Behav. Neurosci. 125, 20–28 (2011).

19. D. G. Gee et al, Early developmental emergence of human amygdala- prefrontal connectivity after maternal deprivation. Proc. Natl. Acad. Sci. U. S. A. 110 (2013), doi:10.1073/pnas. 1307893110.

20. P. Janata, The Neural Architecture of Music-Evoked Autobiographical Memories. Cereb. Cortex. 19, 2579–2594 (2009).

21. S. Koelsch, Towards a neural basis of music-evoked emotions. Trends Cogn. Sci. 14, 131–7(2010).

22. J. Plailly, B. Tillmann, J.-P. Royet, The Feeling of Familiarity of Music and Odors: The Same Neural Signature? Cereb. Cortex. 17, 2650–2658 (2007).

23. K. E. Loveland, D. Smeesters, N. Mandel, Still Preoccupied with 1995: The Need to Belong and Preference for Nostalgic Products. J. Consum. Res. 37, 393—408 (2010).

24. F. S. Barrett, P. Janata, Neural responses to nostalgia-evoking music modeled by elements of dynamic musical structure and individual differences in affective traits. Neuropsychologia. 91, 234–246 (2016).

## Supplemental References

1. J. Talairach, P. Tournoux, Co-planar stereotaxic atlas of the human brain. 3- Dimensional proportional system: an approach to cerebral imaging (1988).

2. R. W. Cox, AFNI: Software for Analysis and Visualization of Functional Magnetic Resonance Neuroimages. Comput. Biomed. Res. 29, 162–173 (1996).

3. K. R. A. Van Dijk et al., Intrinsic functional connectivity as a tool for human connectomics: theory, properties, and optimization. J. Neurophysiol. 103, 297–321 (2010).

4. C.-G. Yan et al., A Comprehensive Assessment of Regional Variation in the Impact of Head Micromovements on Functional Connectomics. NeuroImage. (2013), doi:10.1016/j.neuroimage.2013.03.004.

5. D. G. Gee et al, A developmental shift from positive to negative connectivity in human amygdala-prefrontal circuitry. J. Neurosci. 33, 4584–93 (2013).

6. K. Friston et al, Psychophysiological and Modulatory Interactions in Neuroimaging. NeuroImage. 6, 218–229 (1997).

7. K. J. Preacher, A. F. Hayes, SPSS and SAS procedures for estimating indirect effects in simple mediation models. Behav. Res. Methods, Instruments, Comput. 36, 717–731 (2004).

8. P. Janata, The Neural Architecture of Music-Evoked Autobiographical Memories. Cereb. Cortex. 19, 2579–2594 (2009).

9. S. Koelsch, Towards a neural basis of music-evoked emotions. Trends Cogn. Sci. 14, 131–7 (2010).

10. S. Koelsch et al., The roles of superficial amygdala and auditory cortex in music-evoked fear and joy. NeuroImage. 81, 49–60 (2013).

11. R. W. Wilkins, D. A. Hodges, P. J. Laurienti, M. Steen, J. H. Burdette, Network Science and the Effects of Music Preference on Functional Brain Connectivity: From Beethoven to Eminem. Sci. Rep. 4, 6130 (2015).

12. A. J. Blood, R. J. Zatorre, Intensely pleasurable responses to music correlate with activity in brain regions implicated in reward and emotion. Proc. Natl. Acad. Sci. U. S. A. 98, 11818–23 (2001).

13. S. Koelsch, T. Fritz, D. Yves Cramon, K. Mü ller, A. D. Friederici, Investigating Emotion With Music: An fMRI Study. Hum Brain Mapp. 27, 239–250 (2006).

14. A. C. Tziortzi et al., Imaging dopamine receptors in humans with [11C]-(+)- PHNO: Dissection of D3 signal and anatomy. NeuroImage. 54, 264–277 (2011).

15. P. Morosan et al., Human Primary Auditory Cortex: Cytoarchitectonic Subdivisions and Mapping into a Spatial Reference System. NeuroImage. 13, 684–701 (2001).

16. J. Plailly, B. Tillmann, J.-P. Royet, The Feeling of Familiarity of Music and Odors: The Same Neural Signature? Cereb. Cortex. 17, 2650–2658 (2007).

17. A. J. Blood, R. J. Zatorre, Intensely pleasurable responses to music correlate with activity in brain regions implicated in reward and emotion. Proc. Natl. Acad. Sci. U. S. A. 98, 11818–23 (2001).

18. A. J. Blood, R. J. Zatorre, P. Bermudez, A. C. Evans, Emotional responses to pleasant and unpleasant music correlate with activity in paralimbic brain regions. Nat. Neurosci. 2, 382–387 (1999).

19. S. Brown, C. J. Michael Martinez, L. M. Parsons, Passive music listening spontaneously engages limbic and paralimbic systems (available at https://mimm.mcmaster.ca/publications/pdfs/neuroreport.pdf).

